# Modelling sexual violence in male rats: The sexual aggression test (SxAT)

**DOI:** 10.1101/2021.10.06.463346

**Authors:** Vinícius E. de Moura Oliveira, Trynke R. de Jong, Inga D. Neumann

## Abstract

Sexual assault and rape are crimes that impact victims worldwide. Although the psychosocial and eco-evolutionary factors associated with this antisocial behavior have repeatedly been studied, the underlying neurobiological mechanisms are still unknown mainly due to the lack of an appropriate animal model of sexual aggression (SxA). Here, we established a novel paradigm to provoke and subsequently assess SxA in adult male Wistar rats: the sexual aggression test (SxAT). Briefly, male Wistar rats are sexually aroused by a receptive female, which is exchanged by a non-receptive female immediately after the first intromission. This protocol elicits forced mounting (FM) and aggressive behavior (AB) towards the non-receptive female to different degrees, which can be scored. In a series of experiments we can show that SxA behavior is a relatively stable trait in rats and correlates positively with sexual motivation. Rats with innate abnormal anxiety and aggressive behavior also show abnormal SxA behavior. In addition, central infusion of oxytocin moderately inhibits AB, but increases FM. Finally, we identified the agranular insular cortex to be specifically activated by SxA, and inhibition of this region mildly decreased AB during the SxAT.

Altogether, the SxAT is a paradigm that can be readily implemented in behavioral laboratories as a valuable tool to find answers regarding the biological mechanisms underlying SxA in humans, as well as social decision-making in general.

## 1. INTRODUCTION

According to the World Health Organization (WHO), worldwide, about 35% of women have experienced physical and/or sexual violence in their lifetime^1^, which is even likely to be an underestimation^2^. Rape and sexual assault have severe physical and mental consequences including post-traumatic stress disorder^3,4^. Thus, effective prevention of sexual violence is highly warranted, but depends largely on the identification of factors that facilitate or inhibit a person’s tendency to repeatedly assault or rape. Unfortunately, current risk assessments and therapies aimed to prevent recidivism of sex offenders lack adequate efficacy^5–7^.

Traditionally, the motivating factors contributing to (male-to-female) sexual aggression (SxA) have been studied in two relatively separated scientific worlds. On the one hand, ecologists and evolutionary biologists have shown that sexual conflict is common in many animal species and driven by the evolutionary forces of reproductive success and failure^8–10^. On the other, sociologists, psychologists, and psychiatrists have used questionnaires and diagnostic instruments to formulate theoretical frameworks modeling the important roles of culture, childhood experiences, and personality traits in the human tendency to rape^11,12^.

While it is certainly important to understand the evolution of SxA throughout the animal kingdom and rape culture in humans, empirical data on genetic, hormonal, and neuroanatomical factors underlying an individual’s tendency to display SxA are essential in order to seek effective therapies. Unfortunately, there are only anecdotal descriptions of biological factors identified in (violent) sex offenders. Thus, white matter abnormalities in cortical and subcortical brain areas associated with moral decision-making, reward-processing, sexual arousal, and aggression^13,14^, and increased testosterone or gonadotrophic hormones levels moderately predicting recidivism and hostility, have been described ^15,16^

Detailed investigations into the neurobiology of sexual violence can only be promoted using a validated animal model of SxA under controlled laboratory conditions, involving rodent cohorts with relatively homogeneous sexual behavior and increased SxA. Moreover, only a rodent model of SxA allows experimental, including neuropharmacological interventions, to reveal neurobiological underpinnings of sexual violence. Rodent models have been successfully used for decades to investigate the neuronal mechanisms of various innate occurring or dysfunctional social behaviors including social decision-making, empathy, consolation, social reciprocity, social fear and social avoidance, impaired maternal behavior, and extremes in male or female aggression [reviewed in^17,18^]. Aggression in a sexual context is notably an exception to this list.

Therefore, we aimed to establish a robust rat model of male SxA that could be easily implemented in other behavioral laboratories. Appreciating the enormous diversity of potential genetic and developmental origins, and neurobiological mechanisms of SxA in humans –we aimed to establish a model with high face validity and focused on basic aspects of human rape and sexual assault, i.e. forced sexual acts and aggressive behavior towards a non-willing and defensive female in the absence of sexual compliance.

In the process of establishing the first rat model of SxA, we first assessed the frequency and behavioral variability of forced mounting (FM) and aggressive behavior (AB) towards non-receptive females in sexually aroused male Wistar rats. Secondly, as personality traits^11,14^ have been linked with sexual violence, we investigated, whether innate differences in anxiety and social behaviors affect the display of SxA using rats selectively bred for high (HAB) versus low (LAB) anxiety-related behavior^19,20^. For further validation of the SxA model, we applied intraperitoneal (i.p.) ethanol, known to play a role in human rape and assault^21^. We also tested two pharmacological treatments hypothesized to affect SxA, i.e. intracerebroventricular (i.c.v.) infusion of oxytocin (OXT) or arginine vasopressin (AVP), both known to affect pro-social interactions, aggression, and sexual behavior in male rats^18,21–29^. Finally, we analyzed the patterns of neuronal activity in selected brain areas in response to either sexual aggression, consensual mating, or territorial aggression. Based on these results we functionally inhibited the posterior agranular insular cortex (pAIC) to link neuronal activity and behavior.

## 2. MATERIALS & METHODS

### 2.1. Ethics Statement

Experiments were approved by the Committee on Animal Health and Care of the Government of the Unterfranken (2532-2-443) and followed the Guide for the Care and Use of Laboratory Animals produced by the National Institute of Health.

### 2.2. Animals

Experiments were carried out in adult (12-15 weeks) male Wistar rats, which were bred in the animal facilities of the University of Regensburg, Germany, and either non-selected (NAB), or selectively bred for high (HAB) or low (LAB) anxiety-related behavior^19,20^. As stimulus animals during the various behavioral tests young (10-12 weeks) adult female or male NAB Wistar rats, either bred at the animal facilities of the University of Regensburg or obtained from Charles Rivers Laboratories (Sulzfeld, Germany), were used. All rats were kept under controlled laboratory conditions (12:12h light/dark cycle; lights off at 11:00h, 21±1**°**C, 60±5% humidity, standard rat nutrition (RM/H, Ssniff Spezialdiäten GmbH, Soest, Germany) and water *ad libitum*) and housed in groups of four in standard rat cages (55 × 35 × 20 cm) with sawdust bedding until the start of the experimental procedures. All behavioral tests were performed in the first half of the dark phase, between 12:00h and 17:00h. Additionally, 48 hrs prior to the behavioral procedures (except the elevated plus-maze) the experimental males were single-housed in experimental observation cages (home cage, 40×24×35 cm, plexiglas walls).

### 2.3. Behavioral Tests

#### 2.3.1. Sexual Aggression Test (SxAT)

We established the SxAT to initially provoke SxA and to assess the individual degree of SxA. The SxAT consisted of two distinct, consecutively performed parts: an instigation and a frustration phase. Prior to the SxAT, stimulus females were briefly exposed to a non-experimental sexually experienced male Wistar rat and classified as sexually receptive or non-receptive based on the presence or absence of hopping, darting, and lordosis. In the first phase, sexual instigation was achieved by introducing a receptive female into the male’s homecage; the latency to mount the receptive female (with or without intromission) was measured as copulation latency (CL).

As soon as the male performed either 3 mounts or 1 intromission (whatever occurred first), or until 5 min elapsed without any mount or intromission, the receptive female was replaced by a non-receptive female, thus starting the frustration phase. During the following 10 min, all signs of SxA, i.e. the frequency of FM (the number of attempts to mount the non-receptive female) and AB (the number of aggressive behaviors toward the female). AB comprised of behaviors typically displayed by rats during territorial aggression and scored in the resident-intruder test^30^: “forced grooming” (male aggressively licking the head/neck area of the female), “keep down” (male using his front paws and upper body to force the female to lie on her back), “threat” (male displaying a threatening posture or movement toward the female, including pushing/shoving the female with his head), “lateral threat” (male turning his body sideways and pushing the female into a wall or corner) and “attack” (rapid clinch attack with one or more bites). Non-receptive females were immediately removed from the cage and replaced by another non-receptive stimulus female once they showed any receptive behaviors (hopping, darting, or lordosis),

The CL, and the occurrence of FM and AB were first scored live by a trained observer using a stopwatch and a standardized pen-and-paper score form. All SxAT were also recorded with an infrared video camera (Sumikon, PEARL GmbH, Buggingen, Germany). For pharmacological experiments (Exp 3 and Exp 5), the full spectrum of behaviors throughout the SxAT was scored from video using JWatcher behavioral observation software^31^ in order to detect any changes due to the treatments. Thus, aside from FM and AB, also neutral behaviors (immobility, exploration, eating/drinking, autogrooming) and non-aggressive social interactions (anogenital sniffing, defensive behavior) were scored. From these scores, the percentage of time that an animal spent performing a specific type of behavior was calculated (total duration of behavior / 10 min x 100 %).

#### 2.3.2. Consensual Mating Test (CMT)

To assess ‘normal’ sexual behavior of experimental males, the consensual mating test (CMT) was performed as described^32^. A receptive female was placed in the home cage of the experimental male, i.e. in a non-paced mating condition^33^, and free interactions were allowed for 10 min. Mounting, copulation, and ejaculation latencies, as well as respective frequencies, were scored either live or on video recordings using JWatcher software.

#### 2.3.3. Resident Intruder Test (RIT)

To assess intermale territorial aggression, the resident intruder test (RIT) was performed as described^30^. Briefly, a virgin male intruder, weighing approximately 20% less than the experimental resident male, was placed in the home cage of the resident for 10 min. Various social and non-social behaviors of the resident also scored during the SxAT (please see session 2.3.1. for details) were recorded on an infrared video camera and analyzed using JWatcher software.

#### 2.3.4. Elevated Plus-Maze (EPM) Test

For analysis of anxiety-related behavior, rats were tested on the elevated plus-maze (EPM) ^19,20^ as described in detail in the supplementary information.

### 2.4 Stereotaxic Surgery

Stereotaxic surgery was performed as previously described^26,27^, for further information please see supplementary information.

### 2.5 Pharmacological Interventions

#### 2.5.1. I.c.v. OXT or AVP infusion

Synthetic OXT and AVP (Sigma-Aldrich Biochemicals, Munich, Germany) were dissolved in Ringer’s solution (VEH, pH=7.4, B. Braun AG, Melsungen, Germany). OXT was infused i.c.v. at a dose of 1 or 100 ng/5µl^26,27^, and AVP was infused at a dose of 0.1 or 1 ng/5µl^26^ 10 to 15min prior to the SxAT.

#### 2.5.2. I.p. Ethanol Injection

Ethanol (Fortior Primasprit Neutralalkohol, Brüggemann Alkohol, Heilbronn, Germany) was dissolved in saline (VEH) at 15% w/v, and injected i.p. at either 0.5g/kg or 1.5g/kg 10 min prior to the SxAT.

#### 2.5.3. Infusion of Baclofen and Muscimol into the posterior Agranular Insular Cortex (pAIC)

The GABA-A receptor agonist muscimol (Sigma Aldrich GmbH, Munich, Germany) and the GABA-B receptor agonist baclofen (Sigma Aldrich GmbH, Munich, Germany) were dissolved together in saline at a dose of 20ng/μl (muscimol) and 200ng/μl (baclofen), respectively, and 1μl of this cocktail was slowly infused into the left and right posterior agranular insular cortex 5min before the start of the SxAT.

### 2.6 Immunohistochemistry and Microscopical analyses

Immunohistochemistry was performed as previously described^27^; for further information about analysis, protocol and abbreviations please see supplementary information.

### 2.7. Experimental Protocols

#### 2.7.1 Behavioral profiling of SxA (Experiment 1)

In order to assess (i) the general occurrence of SxA, (ii) the individual variability in SxA among male NAB Wistar rats (n=126), and (iii) the correlations between SxA behaviors and sexual motivation in Wistar rats, we analyzed CL, FM and AB during three SxAT performed on three consecutive days, i.e. SxAT-1/2/3 (described in 2.3.1). Note that some of these rats (n=32) underwent the three SxA tests (SxAT-1/2/3) specifically for experiment 1, whereas the other rats (n=94) were exposed to three consecutive SxATs as a ‘training’ before the pharmacological treatments of experiment 3. Of this latter group, 57 rats had an i.c.v. cannula implanted 3 days prior to SxAT-1 and were handled before each SxAT (see description in 2.7.3).

### 2.7.2. SxA in HAB, NAB, and LAB rats (Experiment 2)

To determine whether an innate high or low level of anxiety-related behavior influences SxA, HAB, NAB, and LAB rats (n=10 each) were tested in three SxAT performed on three consecutive days (SxAT-1/2/3). High, normal, and low anxiety-related behavior was confirmed on the EPM (described in 2.3.4) either one week before (HAB/LAB) or after (NAB) testing in the SxAT.

The standard SxAT paradigm (described in 2.3.1.) had to be modified for this particular experiment to protect stimulus females against the excessive aggression (i.e. lateral threats and attacks) displayed by the LAB and (to a lesser extent) HAB rats, confirming prior findings^34,35^ Therefore, the SxAT were curtailed to 5 min rather than 10, and the instigation phase was skipped in SxAT-2 (i.e. only the frustration phase was performed).

Excessive aggression in a SxAT was analyzed as [the number of attacks and lateral threats] / [the number of all aggressive behaviors] x 100%, with 0/0 counted as 0.

#### 2.7.3 Pharmacological manipulation of SxA (Experiment 3)

For pharmacological manipulation, rats fitted with an i.c.v. cannula 3 days prior to SxAT-1 (see 2.4) were first trained in three SxAT performed on three consecutive days (SxAT-1/2/3). The behavioral results from these training sessions were added to the analyses in experiment 1. The training phase served the purpose a) to habituate males to the administration procedure, b) to stabilize SxA, c) to exclude animals with consistently low SxA scores from the experiment (n=4), and d) to equally distribute the remaining animals to the experimental groups to achieve balanced average SxA levels. One rat was removed from the experiment due to lethargic behavior in the home cage.

Effects of i..c.v. pharmacological treatments were assessed in SxAT-4, performed one day after SxAT-3. For experiment 3a, rats were infused with either VEH (n=9), 1ng/5µl of OXT (OXT-1, n=10) or 100ng/5 µl of OXT (OXT-100, n=9) 15 min prior to SxAT-4 (see 2.5.1).

For experiment 3b, rats were infused with either VEH (n = 9), 0.1ng/5µl of AVP (AVP-0.1, n=8) or 1ng of AVP/5µl (AVP-1, n=10) 10 min prior to SxAT-4 (see 2.5.1).

For experiment 3c, rats were i.p. injected with either saline (VEH, n=11), 0.5mg/kg ethanol (ETH-0.5, n=11) or 1.5mg/kg (ETH-1.5, n = 12) 10 min prior to SxAT-4 (see 2.5.2).

In experiments 3a, b, and c, exposure to the SxAT-4 was followed immediately by a 5-min CMT (see 2.3.2) to assess, whether the pharmacological treatments had affected general sexual motivation. Animals, who responded to treatment with abnormal behavior (for example extensive immobility or excessive auto grooming) were removed from statistical analyses (OXT-1: n=1; OXT-100: n=2; AVP-1: n=1; ETH-1.5: n=4).

#### 2.7.4 Neuronal activation in response to SxA (Experiment 4)

To monitor the neuronal activity in response to sexual aggression within relevant brain regions, male Wistar rats (n=36) were exposed to three CMTs performed on three consecutive days to a) habituate them to the behavioral test set-up, b) to gain an equal level of sexual experience, and c) to facilitate interactions with stimulus animals on the final experimental day 4.

On day 4, males were equally distributed to four experimental groups, receiving either no stimulus (control group), 10-min exposure to a non-receptive female (SxAT), 10-min exposure to a receptive female (CMT), or 10-min exposure to a smaller male intruder (RI test). In the control group, an observer mimicked the placement of a stimulus animal in the cage. Note that the instigation phase was omitted from the SxAT in this experiment to avoid any neuronal activation in response to consensual mating in this group. One hour after exposure to the different test conditions, animals were transcardially perfused, and brains were prepared for immunohistochemistry (see 2.6).

Five males were excluded from analysis due to either low overall sexual motivation during training or to suboptimal perfusion/brain tissue processing, resulting in the following group sizes: n=4 (control), n=12 (SxAT), n=8 (CMT), n=7 (RIT). The following behaviors were selected for correlation analyses: number of FM (SxAT) and AB (SxAT, RIT), and mounting (CMT).

#### 2.7.5 Effects of pAIC inhibition on SxA (Experiment 5)

To test whether the pAIC – which was found to be particularly activated in response to SxA - is causally linked to the display of SxA behavior, we functionally inhibited the pAIC prior to a SxAT. As explained above (see 2.7.3), rats bilaterally fitted with guide cannulas above the pAIC 5 days prior to training were first trained in three test sessions performed on three consecutive days. To homogenize sexual behavior between subjects, we modified the training paradigm to include two CMTs (CMT-1/2) followed by one SxAT (SxAT-1). The experimental phase consisted of three consecutive daily SxAT (SxAT-2/3/4), in which rats received either a muscimol-baclofen cocktail or saline (see 2.5.3 for details) in a cross-over within-subjects design 5 min prior to either SxAT-2 or SxAT-4. SxAT-3 served as a wash-out test. Eleven rats were excluded from analyses due to either low sexual motivation during the two CMTs (n=7) or misplaced guide cannula (n=4).

### 2.8. Statistics

All statistics were performed using SPSS version 23 (IBM).

For experiment 1, behavioral changes over time were analyzed using a 2-way mixed ANOVA with time (i.e. three consecutive daily SxAT) as within-subjects factor and surgery (none vs. i.c.v. cannula placement) as between-subjects factor. In addition, Pearson’s correlation analyses were performed to assess the correlations between the various behavioral outcomes. For experiment 2, ANOVA followed by Bonferroni-corrected post-hoc pairwise comparisons were performed to assess behavioral differences in the SxAT between HAB, LAB and NAB rats. For experiment 3, ANOVA followed by Dunnett’s pairwise comparisons of treatments with VEH were performed to assess the effects of i.c.v. OXT, i.c.v. AVP, and i.p. ethanol on behavior in the SxAT. For experiment 4, ANOVA followed by Bonferroni-corrected posthoc pairwise comparisons were performed to assess group differences in Fos-IR. In addition, Pearson’s correlation analyses were performed to assess the correlations between specific behavioral parameters and Fos-IR.

For experiment 5, within-subjects two-tailed Student’s t-tests were used to compare behavioral scores following vehicle versus muscimol-baclofen treatment.

## 3. RESULTS

### 3.1. Experiment 1: Basic behavioral profile and individual differences in SxA

CL significantly declined over the three consecutive SxAT (F[2] = 41.909, p < .001) (Fig. 1A), whereas there were no changes in the frequency of FM (F[2] = .090, p = .914) or AB (F[2] = 1.457, p = .235) (Fig. 1B and 1C, respectively). As none of the behaviors were affected by i.c.v. surgery (F[1] < 1.412, p > .246), behavioral data from intact and cannulated animals were pooled for the correlation analyses.

**Fig. 1.**
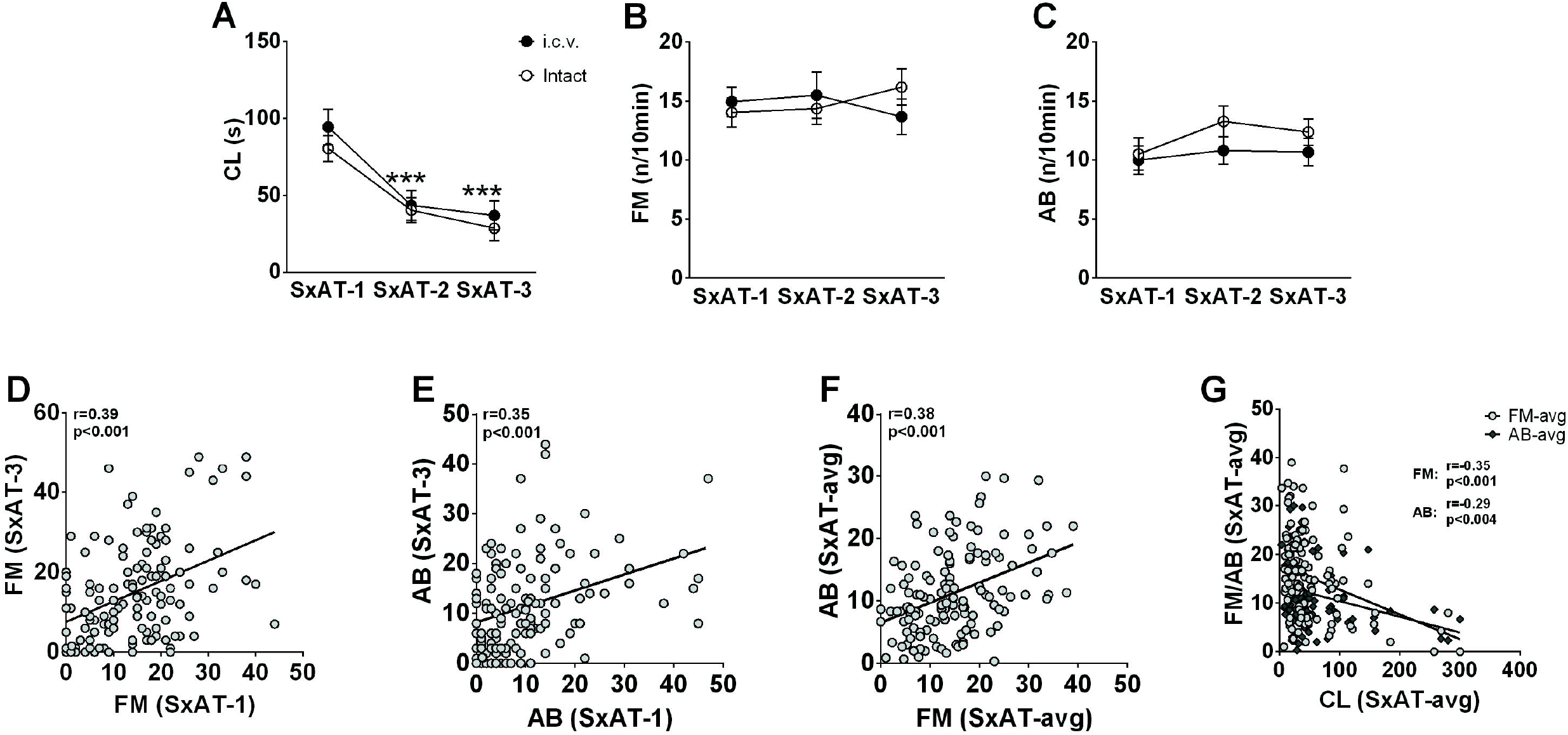
Behavioral profile of adult male Wistar rats in sexual aggression tests (SxAT). (**A**) Latencies to copulate (CL) with a receptive female, (**B**) frequency of forced mounts (FM), and (**C**) frequency of aggressive behavior (AB) of intact (white circles) and i.c.v. cannulated (black circles) males towards a non-receptive female during the SxAT performed on three consecutive days. Data are means +/-S.E.M. **D**: Scatter plot of FM frequencies displayed during SxAT-1 vs. SxAT-3. (**E**) Scatter plot of AB frequencies during SxAT-1 vs. SxAT-3. (**F**): Scatter plot of FM frequencies (averaged over SxAT-1, SxAT-2 and SxAT-3) vs. AB frequencies (averaged over SxAT-1, SxAT-2 and SxAT-3). (**G**) scatter plots of the CL vs. FM (dark grey diamonds) or AB (grey circles) averaged over SxAT-1, SxAT-2 and SxAT-3).

Relevant Pearson’s correlation coefficients among the behavioral parameters in SxAT1-3 are summarized in Table 1. The FM frequency scores were found to significantly correlate with each other in all three training sessions, with the strongest correlation between SxAT-1 and SxAT-3 (Fig. 1D). Also, the aggression scores correlated positively with each other in the three training sessions, with the strongest correlation between SxAT-1 and SxAT-3 (Fig. 1E). The FM and AB frequencies correlated positively and significantly within each training session, resulting in a significant correlation between FM and AB scores averaged over SxAT-1/2/3 (Fig. 1F). Moreover, a significant negative correlation between averaged CL and FM as well as AB scores was found (Fig. 1G).

**Table 1.** Overview of Pearson’s correlation coefficients between the different behaviors (CL, FM, AB) in each of the three consecutive daily SxAT, or the average over all three SxAT. Significant correlations in bold.

| Correlation |  | r | p |
| --- | --- | --- | --- |
| FM | SxAT-1 vs. SxAT-2 | **.326** | **< .001** |
|  | SxAT-2 vs. SxAT-3 | **.280** | **.001** |
|  | SxAT-1 vs. SxAT-3 | **.399** | **< .001** |
| AB | SxAT-1 vs. SxAT-2 | .085 | .342 |
|  | SxAT-2 vs. SxAT-3 | **.271** | **.002** |
|  | SxAT-1 vs. SxAT-3 | **.351** | **< .001** |
| SxAT-1 | FM vs. AB | **.241** | **.007** |
|  | FM vs. CL | **-.296** | **.002** |
|  | AB vs. CL | -.199 | .037 |
| SxAT-2 | FM vs. AB | **.266** | **.003** |
|  | FM vs. CL | **-.302** | **.001** |
|  | AB vs. CL | -.235 | .013 |
| SxAT-3 | FM vs. AB | **.297** | **.001** |
|  | FM vs. CL | **-.276** | **.004** |
|  | AB vs. CL | -.135 | .163 |
| SxAT-avg | FM vs. AB | **.386** | **< .001** |
|  | FM vs. CL | **-.351** | **< .001** |
|  | AB vs. CL | **-.298** | **< .001** |

### 3.2. Experiment 2: SxA in HAB, NAB, and LAB rats

Statistics for experiment 2 are summarized in Table 2.

**Table 2:**
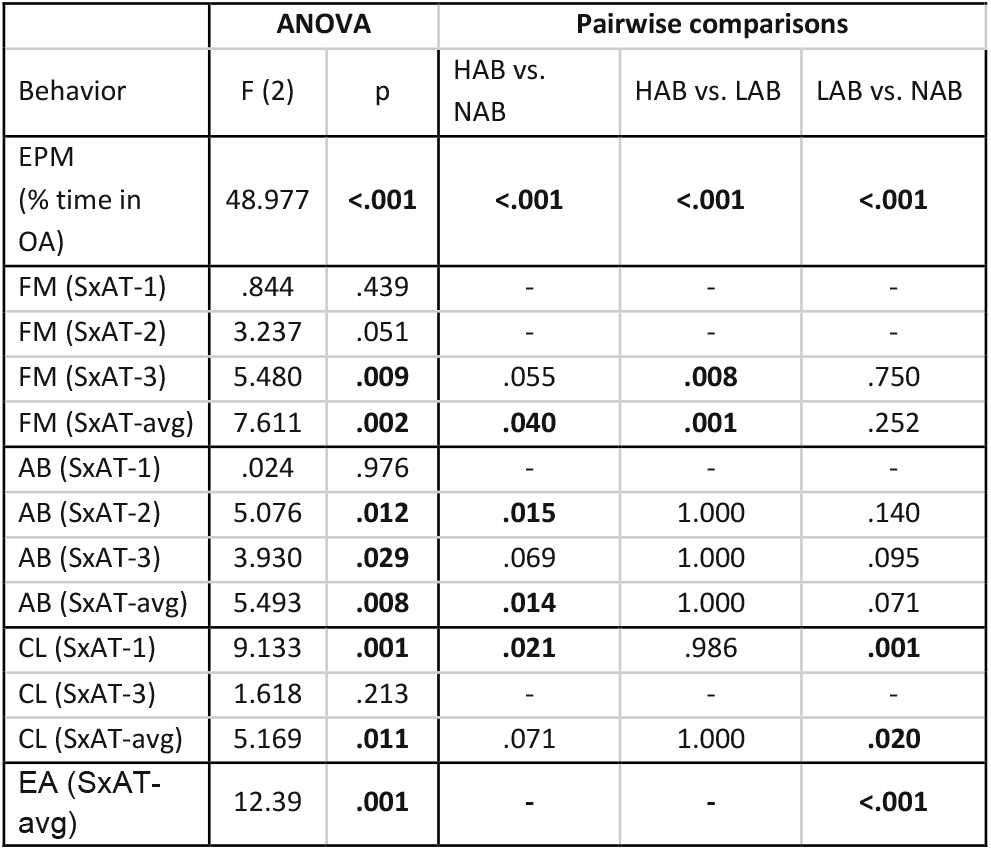
Overview of ANOVA and, if appropriate, post-hoc pairwise comparisons between rats selectively bred for high (HAB) vs low (LAB) anxiety behavior or not selectively bred (NAB) of their behaviors in a 5-min EPM or 10-min SxAT. Significant differences in bold.

HAB, LAB, and NAB rats differed in their levels of anxiety-related behavior on the EPM with HAB rats being the most anxious in the EPM (% of time in open arms: 5.07 ± 1.17), followed by NAB rats (32.35 ± 3.64) and LAB rats (60.33 ± 3.78).

In the SxAT, HAB rats showed the highest FM score, which differed significantly from LAB rats in SxAT-3 (Fig. 2B) and from both LAB and NAB rats in the average of all SxAT (Fig. 2D). HAB rats also showed the highest AB score, which differed significantly from NAB, but not LAB, rats in SxAT-2 (Fig. 2B) and in the average of all SxAT (Fig. 2D). NAB rats showed the longest CL, which differed significantly from both HAB and LAB rats in SxAT-1 (Fig. 2A) and from LAB rats (but not HAB rats) in the average of SxAT-1 and SxAT-3 (Fig. 2D). LAB rats showed more excessive aggression than NAB, but not HAB rats (Fig. 2D).

**Fig. 2.**
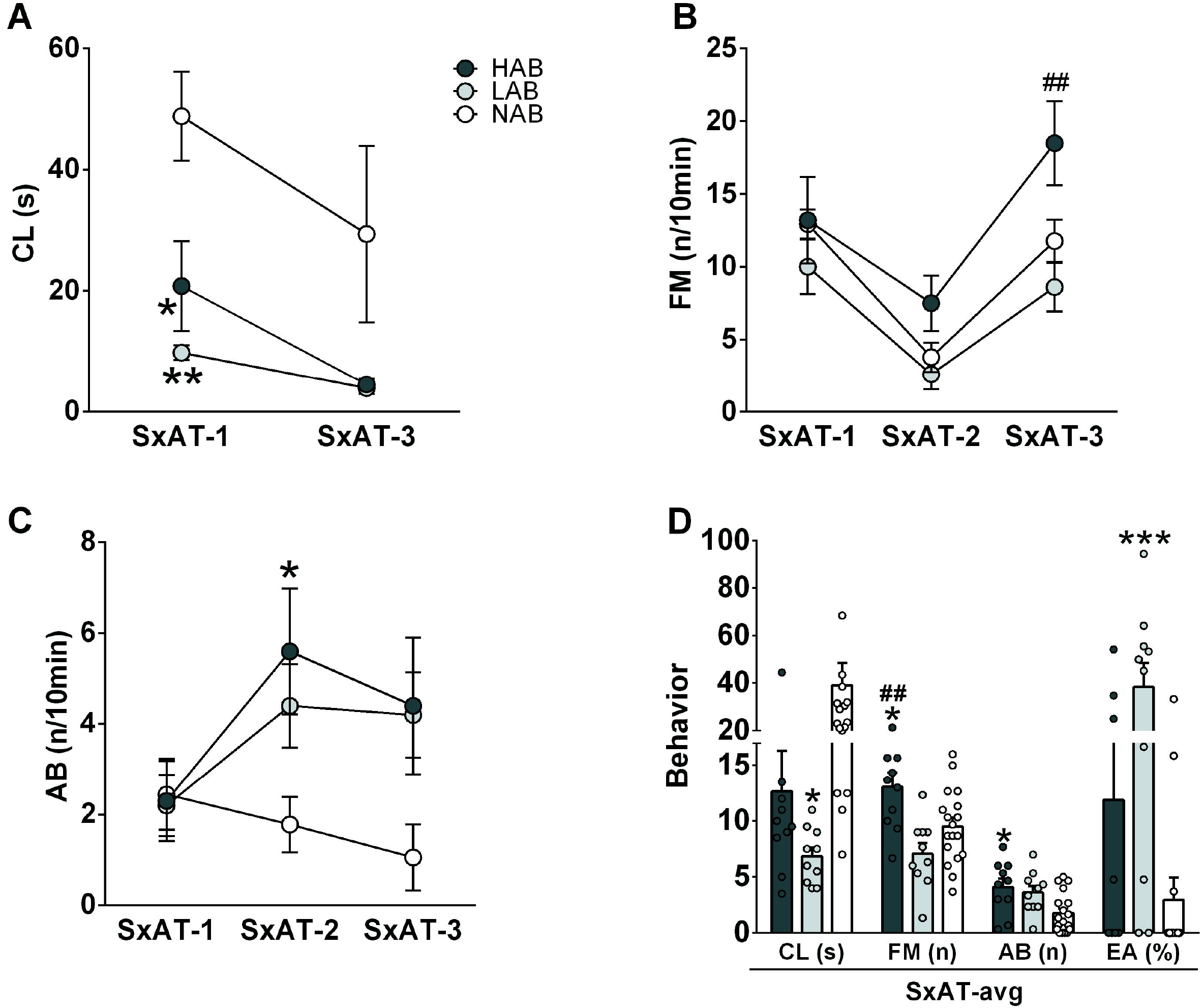
Behavioral profile of adult male HAB (dark-grey circles and bars), LAB (light-grey) and NAB (white) rats in three consecutive daily sexual aggression tests (SxAT-1/2/3). **A**: Latency to copulate (CL) with a receptive female rat in SxAT-1 and SxAT-3. **B**: Frequency of forced mounts (FM), and **C**: Frequency of aggressive behaviors (AB) towards a non-receptive female in SxAT-1/2/3. **D**: CL, FM, AB, and excessive aggression (EA) averaged over SxAT1/2/3. Data are presented as mean +/-s.e.m. * p<0.05, ** p<0.01, *** p<0.001 vs. NAB. ## p<0.01HABvs LAB

### 3.3. Experiment 3: Pharmacological manipulation of SxA

Statistics for experiment 3 are summarized in Table 3.

**Table 3:** Overview of ANOVA and, if appropriate, post-hoc pairwise comparisons of behaviors in a 10-min SxAT in response to i.c.v. infusions of OXT or AVP vs. VEH, or IP injections of ETH vs. VEH. Significant differences in bold.

| Behavior (% of time) | ANOVA |  | Pairwise comparisons |  |
| --- | --- | --- | --- | --- |
|  | F (2) | p | VEH vs. OXT-1 | VEH vs. OXT-100 |
| FM | 5.054 | **.016** | **.029** | .893 |
| AB | 4.139 | **.031** | .629 | **.019** |
| neutral | 1.257 | .305 | - | - |
| social | 0.654 | .530 | - | - |
| Behavior (% of time) | F (2) | p | VEH vs. AVP-0.1 | VEH vs. AVP-1 |
| FM | 0.549 | .585 | - | - |
| AB | 0.321 | .726 | - | - |
| neutral | 0.529 | .596 | - | - |
| social | 0.824 | .451 | - | - |
| Behavior (% of time) | F (2) | p | VEH vs. ETH-0.5 | VEH vs. ETH-1.5 |
| FM | 0.169 | .845 | - | - |
| AB | 2.172 | .134 | - | - |
| neutral | 3.891 | **.033** | .070 | **.030** |
| social | 8.253 | **.002** | **.01** | **.001** |

OXT-treatment: There were group differences in the percentage of time displaying FM and AB, but not in the percentage of time displaying neutral or non-aggressive social behaviors during SxAT-4 between i.c.v. OXT- and vehicle-treated rats. Specifically, OXT-1 (but not OXT-100) treatment significantly increased the percentage of FM compared to vehicle, whereas OXT-100 (but not OXT-1) treatment reduced the percentage of AB compared to vehicle (Fig. 3A). Moreover, there were no group differences in CM assessed in the 5-min CM test.

**Fig. 3.**
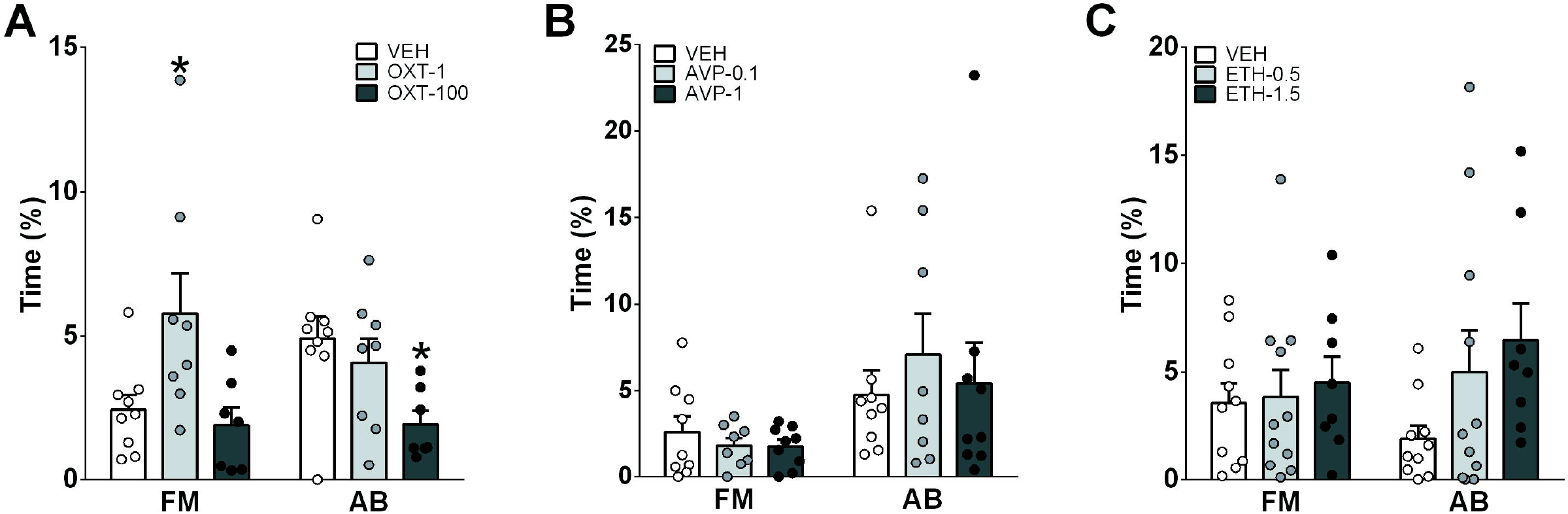
Effects of pharmacological interventions on sexual aggression, i.e., the percentage of time displaying forced mounting (FM) and aggressive behavior (AB) of adult male Wistar rats in a 10-min sexual aggression test (SxAT) following **A)** i.c.v. infusion with OXT (1 or 100ng/5μl Ringer), **B)** i.c.v. infusion with AVP (0.1 or 1ng/5μl Ringer), or **C)** i.p. injection with either ethanol (0.5 or 1.5mg/kg ethanol), or vehicle (VEH, saline). Data are mean + s.e.m. * p< 0.05 vs VEH

AVP-treatment: There were no group differences in any of the behavioral categories measured in the SxAT or CM test between i.c.v. AVP- and VEH-treated rats (Fig. 3B).

Ethanol-treatment: There were no differences in the percentage of FM or AB between ethanol- and vehicle-treated rats (Fig. 3C). However, ethanol-treatment affected the percentage of neutral and non-aggressive social behaviors with ETH-1.5, but not ETH-0.5, treatment significantly increasing neutral behaviors whereas both ETH-0.5 and ETH-1.5 reduced non-aggressive social behaviors (data not shown).

### 3.4. Experiment 4: Neuronal activation in response to SxA

Statistics for experiment 4 are summarized in Table 4.

**Table 4:**
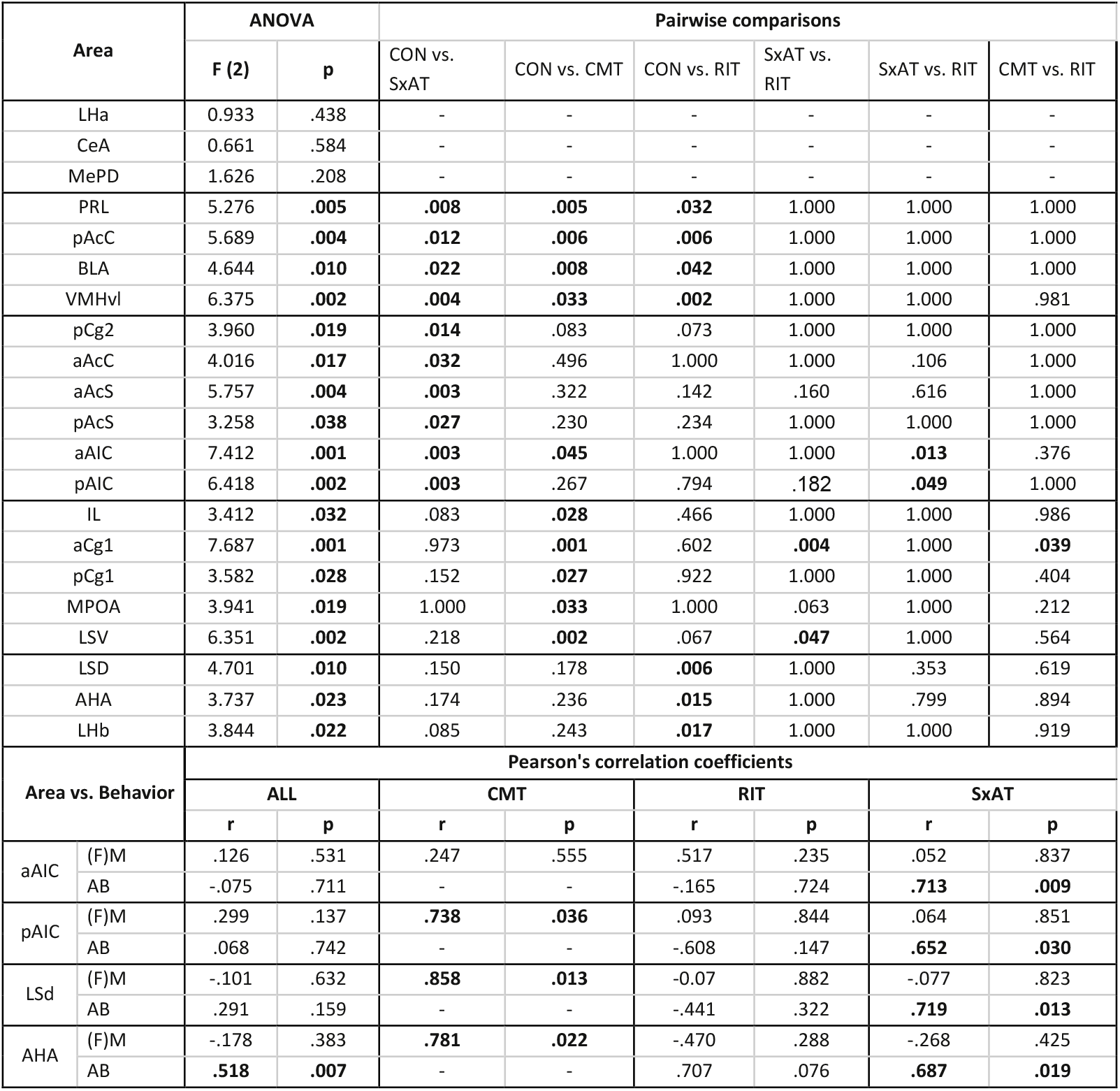
Overview of ANOVA and, if appropriate, post-hoc pairwise comparisons of Fos-IR in various brain regions in response to the experimental conditions (Control, SxAT, CMT, RIT), and Pearson’s correlation coefficients between Fos-IR and individual behaviors. Significant differences and correlations in bold.

ANOVA analysis revealed group differences in Fos-IR in almost all of the analyzed areas, except for the LHA, CeA, and MePD. In areas with group differences, four general statistical patterns could be observed (Fig. 4). Pattern 1 was a similar increase in Fos-IR in response to all behavioral tests compared to control conditions, which was seen in the PRL, pAcC, BLA and VMHvl (Fig. 4A). Pattern 2 was an increase in Fos-IR specifically in response to the CMT compared to control conditions, which was seen in the IL, aCg1, pCg1, MPOA and LSV (Fig. 4B). In the aAIC, Fos-IR was also slightly increased in response to the CMT compared to control conditions. Pattern 3, was an increase in Fos-IR specifically in response to the SxAT compared to control conditions, which was seen in the pCg2, aAcC, aAcS, pAcS, aAIC and pAIC (Fig. 4C). Pattern 4 was an increase in Fos-IR specifically in response to the RIT compared to control conditions, which was seen in the LSD, AHA and LHb (Fig. 4D).

**Fig. 4.**
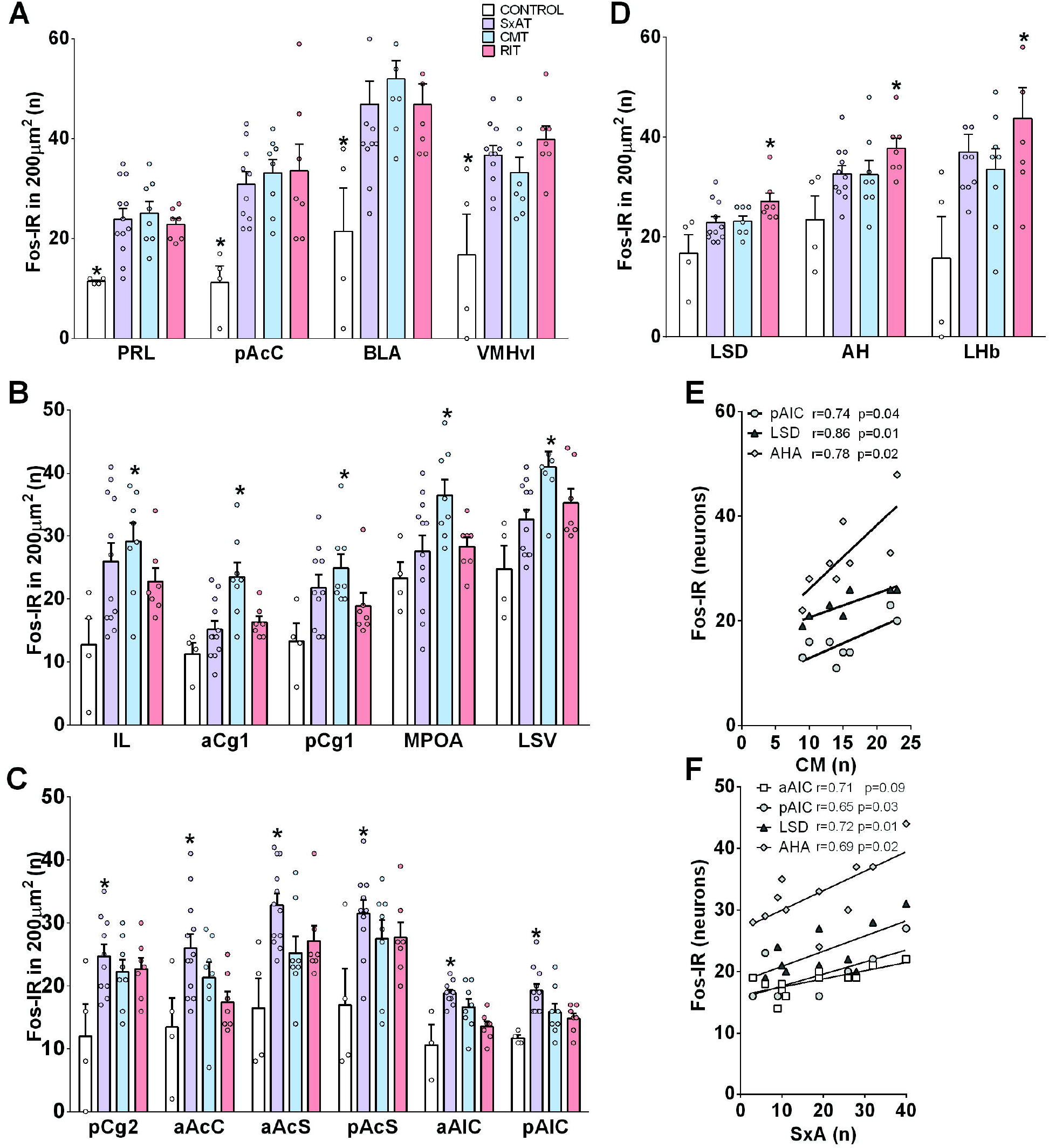
Neuronal activity in selected brain regions indicated by the number of Fos-IR neurons in response to 10 min of rest (CONTROL, white bars) or of exposure to the sexual aggression test (SxAT, purple bars), consensual mating test (CMT, blue bars) or resident-intruder test (RIT, red bars). Brain areas are grouped according to activation patterns: **A)** regions activated in all conditions, **B)** regions activated exclusively by CMT, **C)** regions activated only during the SxAT, and **D)** regions activated only during the RIT. Data are presented as mean + s.e.m.; * p<0.05 vs. CONTROL. **E-F**: Scatter plots of the number of consensual mounts (CM) in the CMT (**E**), or of aggressive behavior (**F**) in the SxAT versus the number of Fos-IR in the aAIC, pAIC, LSD, and AHA. Abbreviations: PRL (pre-limbic cortex); IL (infra-limbic cortex); aCg1 (anterior cingulate cortex 1); pCg1 and pCg2 (posterior cingulate cortex 1 and 2, respectively); aAIC and pAIC (anterior and posterior agranular insular cortex, respectively); aAcC and pAcC (anterior and posterior nucleus accumbens core, respectively); aAcS and pAcS (anterior and posterior nucleus accumbens shell); BLA (basolateral amygdala); CeA (central amygdala); MeApd (posterodorsal medial amygdala); LSD (dorsal lateral septum); LSV (ventral lateral septum); AH (anterior hypothalamic attack area); LHA (lateral hypothalamic attack area); LHb (lateral habenula); MPOA (medial preoptic area); VMHvl (ventrolateral ventromedial hypothalamus).

**Fig. 5.**
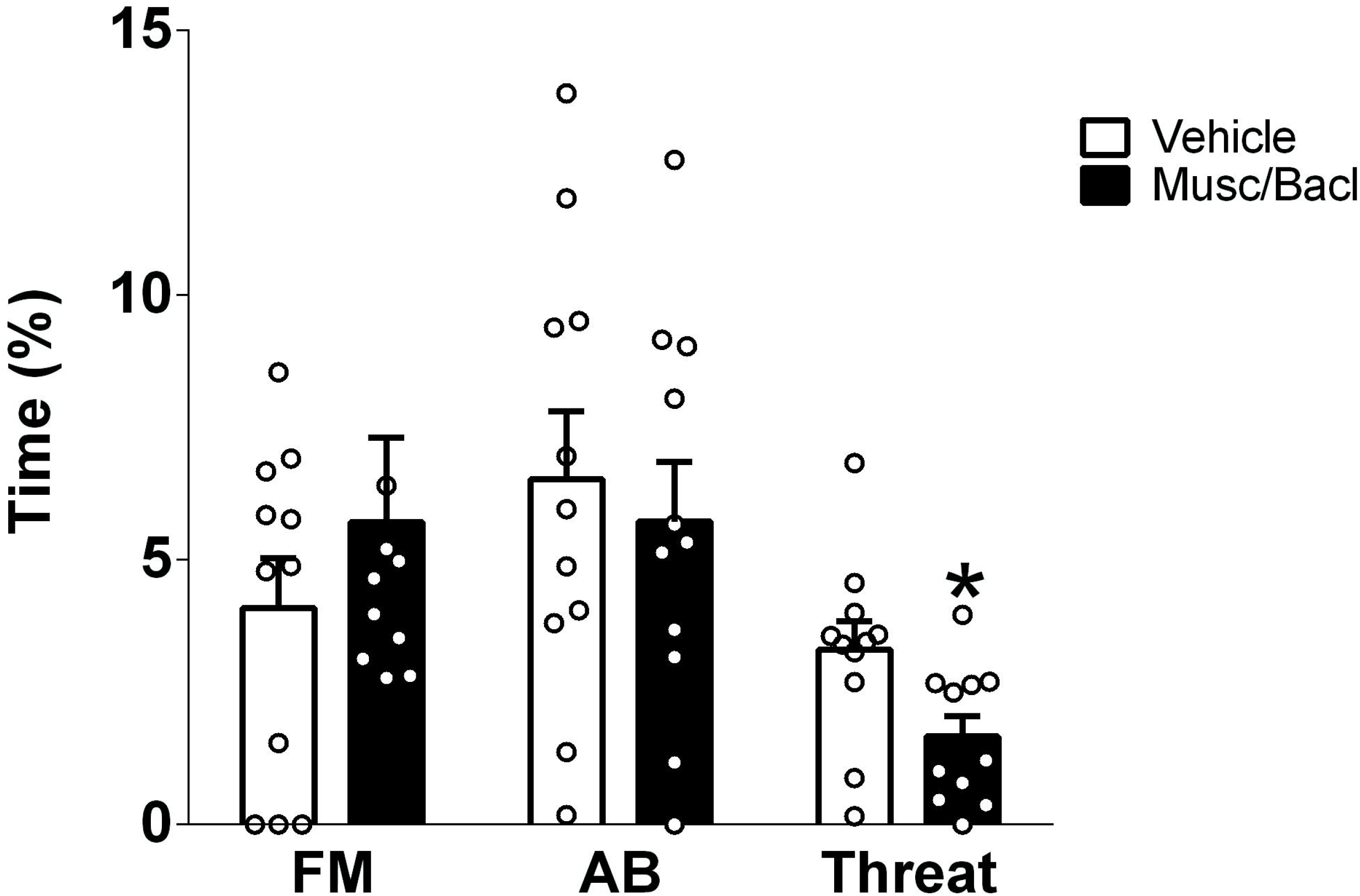
Effects of pharmacological inhibition of the posterior agranular insular cortex (pAIC) on sexual aggression (SxA) indicated by the percentage time displaying forced mounting (FM), and on general aggressive behavior (AB), and threat in particular. Adult male Wistar rats were bilaterally infused with a cocktail of 20ng/μl muscimol and 200ng/μl baclofen or vehicle into the pAIC 5 min prior to exposure to the SxAT. Data are presented as mean + s.e.m.; * p<0.05.

Rats displayed a considerable behavioral individual variability in the various tests, allowing the analysis of correlations between certain behaviors and Fos-IR. We chose consensual mounts (CMT), forced mounts (SxAT, RIT), sexual aggression (SxAT) and territorial aggression (RI test) as the most relevant behaviors. Pearson’s correlation analyses revealed that mounts (in all behavioral tests combined) or FM (in either the SxAT or RIT) did not correlate with Fos-IR in any of the analyzed brain areas, whereas CM (in the CMT) correlated positively with Fos-IR in the pAIC, LSD and AHA (Fig. 4E). Pearson’s correlation analyses furthermore revealed that aggression (in all behavioral tests combined) correlated positively with Fos-IR in the AHA, whereas specific territorial aggression (RIT) showed no correlations with Fos-IR in any of the analyzed brain areas. Importantly, overall SxA (SxAT) correlated positively with Fos-IR in the aAIC, pAIC, LSD and AHA (Fig. 4F).

### 3.5. Experiment 5: Effects of pAIC-inhibition on SxA

Statistics for experiment 5 are summarized in Table 5.

**Table 5:** Detailed behavioral analyses showing the percentage of time spent on: sexual aggression (SxA): general aggression (AB) and forced mounts (FM), highlighting the threat; neutral and social behaviors after the injection of either vehicle or the cocktail of muscimol and baclofen into the pAIC

| Behavior | VEH | Musc/Bacl | p value |
| --- | --- | --- | --- |
| FM (%) | 3.9±0.9 | 4.4±1 | .628 |
| AB (%) | 6.5±1.3 | 5.7±3.7 | .745 |
| threat (%) | 3.3±0.5 | 1.7±0.4 | **.039** |
| **neutral (%)** | 65.9±1.4 | 65.01±3.4 | .765 |
| **social (%)** | 17±2.9 | 13.1±2.1 | .291 |

Inhibition of pAIC significantly decreased the percentage of time spent on threat behavior of the male towards the female during the SxAT (two-tailed Student’s t-test t_(10)_=2.36, p=0.039), whereas the percentage of time spent in FM and AB remained unchanged. Also, sexual motivation towards the receptive female was not affected by the treatment (not shown).

## 4. DISCUSSION

The experimental results reported in this paper indicate that a simple confrontational behavioral paradigm, i.e., the sexual aggression test (SxAT) in rats can be used as a promising and reliable animal model to study SxA and its neurobiological correlates. This is of high translational relevance, as animal models have proven to be essential instruments in the understanding of the mechanisms and biological background of virtually all human behaviors, with the notable exception of sexual violence. Our results show that SxA, i.e. a combination of forced mounting (FM) and aggressive behavior (AB) in a sexual context are readily displayed by male Wistar rats and can be reliably quantified in the SxAT. Although FM and AB vary considerably between individual rats, and from one test to another, we found overall tendencies when analyzing a large cohort of rats or rats with inbred extremes in anxiety and aggressive behaviors, such as HAB and LAB rats. Moreover, SxA displayed during the SxAT was found to be stable enough to distinguish the effects of pharmacological manipulations. We could further identify specific neuronal activation patterns related to SxA, which could be clearly distinguished from those related to territorial aggression and consensual mating, indicating that the pAIC and the nucleus accumbens are most likely to be implicated in SxA behavior. Indeed, inhibition of the pAIC decreased threat behavior, one of the components of sexual aggression, however other parameters of SxA remained unchanged.

Experiment 1 revealed that SxA is an innate behavior spontaneously displayed by most naive male rats without a learning period, which remains stable over three consecutive SxAT. Thus, rats displaying a high spontaneous propensity for SxA indicated by high FM scores tended to keep this behavior over time, and also showed a higher propensity for general AB, suggesting either a common cause for both behaviors or one behavior stimulating the other. Furthermore, FM was negatively correlated with the latency to copulate with a receptive female in the instigation phase. Combined, these correlations indicate that highly sexual motivated rats (low average CL) show an increased tendency for SxA which might lead to increased general aggression in response to repeated rejection by the female. Conversely, rats with lower sexual motivation (high average CL) tend to show lower SxA (both FM and AB). Notably, not all males followed this narrative: in some cases, an extremely passive or aggressive response from the non-receptive female inhibited the SxA behavior of a highly motivated male (data not shown). In other cases, males that were slow to mount receptive females were much more eager to force mounting of non-receptive females. Finally, several males consistently displayed above-average AB combined with below-average FM, and vice versa.

In experiment 2, the conclusions from the first experiment could be confirmed. The increased sexual motivation (i.e., reduced CL) of both HAB and LAB males was accompanied by an increased SxA compared with NAB rats. The association between low CL and high FM and - to a lesser extent - high AB was especially evident in HAB rats, whereas LAB rats showed a more deviant pattern consisting of low CL, average FM and excessive AB (i.e., high level of lateral threats and attacks) toward the females. These results correspond with earlier findings of increased territorial aggression in male HAB and LAB rats, and abnormal aggression (including the attack of female intruders) displayed by LAB rats in the RIT^20,27^. The relevance of these results in rats selectively bred for extremes in emotionality is that increased SxA behavior is not a ‘stand-alone’ deviance, but is part of a complex pattern of abnormal social behaviors of HAB and LAB rats described before.

Experiment 3 was designed as a first approach to pharmacologically manipulating SxA behavior. Based on their proven capacity to promote sexual behavior (OXT^18,24^) and to differentially modulate territorial aggression (OXT and AVP^18,25,26,29,36–39^), and other relevant social behaviors^17,18,40,41^, the OXT and AVP systems were selected as targets. We could show that synthetic OXT at a lower dose increased the occurrence of FM, but reduced AB of male rats towards a non-receptive female intruder. Furthermore, OXT appeared to weaken the correlation between FM and AB by stimulating FM and reducing AB. However, OXT has been applied i.c.v., and our previous data in the context of anxiety, demonstrates that the effects of OXT might vary depending on the site of injection (i.c.v. vs local)^42–44^. Thus, our results can only partly claim a significant role of the OXT system in SxA behavior. Nevertheless, they add an important piece of evidence to the reported anti-aggressive effects of OXT in males^22,25,26,36^, by suggesting that OXT reduces aggression in male residents regardless of the sex of the intruder.

Nevertheless, we suggest implementing the SxAT in pharmacology-based behavioral experiments, similar to tests for sexual behavior or territorial aggression. However, we recommend the following adjustments to SxA experiments to achieve stable and robustly high patterns of SxA prior to neuropharmacological intervention: i) The initial group sizes must be large to enable the selection of individuals with reliable SxA behavior displayed during the full SxA training in order to minimize individual variability. (ii) Also, prolonged social isolation prior to the SxA training experiment may stimulate robust SxA behavior to reach this aim^26^. (iii) A sufficient number of non-receptive stimulus female rats is needed to replace very passive or aggressive individuals. (iv) Extensive and longer post-surgical habituation may avoid stress-induced behavioral inhibition of SxA.

Experiment 4 revealed that exposure to the SxAT in general, and the display of SxA behaviors in particular, induce a specific neuronal activation pattern that only partly overlaps with that induced by consensual mating (CMT) or territorial aggression (RIT). The results implicate a specific role of the cingulate cortex, the nucleus accumbens core and shell, and most prominently the agranular insular cortex in SxA, which show high neuronal activation after exposure to the SxAT. The insular cortex is the core brain area involved in social decision-making, i.e. the facilitation or inhibition of behaviors by positive or negative cues from conspecifics^45,46^. Indeed, exposure to the SxAT can be viewed as a prime example of social decision-making: the clear negative cues from the non-receptive female (kicking, moving the genitals away from the male, audible distress vocalizations) inhibit the ongoing sexual behavior of the male with various degrees of effectiveness depending on (presumably) the male’s sexual motivation and empathic sensitivity toward negative cues, combined with the strength of the cues from the female.

Although neuronal activity data as well as human literature suggest a crucial role of the posterior agranular insular cortex in modulating aggression, sexual behavior, and coercive sexual interactions^45^, we only found mild effects on SxA, i.e. a reduction in threat behavior after pAIC inhibition. This could be explained due to two main reasons. First, similarly to aggression^47^ and mating^48^, SxA is likely to be encoded in complex circuits involving multiple brain regions. Thus, we hypothesize that some aspects of this behavior will be modulated by different networks, perhaps the pAIC is recruited only in the aggression-related pathway, whereas other brain regions such as the nucleus accumbens play a stronger role in the sexual motivation aspects of SxA behavior (FM). Second, local drug infusion will target all neuronal populations of the pAIC, which is known to be a heterogeneous brain region with different layers and presumably various neuronal populations^49,50^. More selective experimental approaches are needed to functionally dissect the neuronal populations within the pAIC, which are involved in the modulation of SxA.

Taken together, the present experiments introduce the SxAT as a protocol for studying sexual aggression. It uses innate behavior, readily displayed and easily reproduced by any lab experienced in rat behavioral analyses. It could be utilized to study the underlying neurobiological mechanisms of sexual aggression in humans, and potentially to test, if certain contexts or treatments can inhibit this antisocial behavior. The SxAT may also be added to the growing arsenal of paradigms to study social decision-making as the test solely relies on the fact that the male actively “decides” to continue his attempts to copulate with a non-receptive female, despite her clear rejective and distressing responses. In future studies, this paradigm could be extremely relevant for understanding behavioral, neurobiological, and hormonal consequences of sexual defeat in females. This would allow scientists to get knowledge about the psychopathological consequences of coercive sex in humans.

## 5. ACKNOWLEDGMENTS

We would like to thank Rodrigue Maloumby, Sarah Funkenhauser, Günes Birdal, and Gagik Yeghiazaryan for their technical help. This work was supported by the EU FP7 Project: Neurobiology and Treatment of Adolescent Female with Conduct Disorder: The Central Role of Emotion Processing Fem-NATCD (602407; I.N.), the German Research Foundation (GRK 2174, NE465/31, NE465/33; I.N.)

## 6. AUTHOR CONTRIBUTIONS

T.d.J. and V.O. performed and analyzed the experiments, prepared the first draft of the manuscript and of all figures. I.D.N. substantially revised the manuscript. T.d.J. conceived the project with critical input of I.D.N.

