## Supplementary information for "Modelling sexual violence in male rats: The sexual aggression test (SxAT)"

Elevated Plus-Maze (EPM) Test

For analysis of anxiety-related behavior, HAB and LAB males were tested on the elevated plus-maze (EPM) for 5 min in the early dark phase (13.00-16.00h) as described[19,20]. The time spent in open and closed arms, and number of entries into closed arms were recorded using a video/computer system (Plus-maze version 2.0; Ernst Fricke). Anxiety-like behavior is displayed as percentage of time spent on the open arms (time on open arms / [time on open + closed arms] x 100%), locomotor activity as number of entries into closed arms.

Stereotaxic Surgery

Animals were anesthetized with isoflurane, (Forene®, Abbott GmbH & Co. KG, Wiesbaden, Germany), and treated i.p. with an analgesic (0.05 mg/kg Buprenorphine, Bayer, Germany) and an antibiotic (Baytril®,10 mg/kg, Bayer Vital GmbH, Leverkusen, Germany), before the guide cannula was stereotaxically implanted. The i.c.v. cannulas (21G, 12mm; Injecta GmbH, Germany) was implanted above the left lateral ventricle (1.0mm caudal to bregma, 1.6mm lateral to midline, 2.0mm beneath the skull surface[34]). The cannulas for local infusion (26G, 12mm; Injecta GmbH, Germany) were implanted bilaterally 2 mm above the pAIC (1.6mm rostral, 4.5mm lateral, 6.0mm deep). Cannulas were fixed to the skull with two jeweler's screws and dental cement (Kallocryl, Speiko-Dr. Speier GmbH, Muenster, Germany), and closed by a stainless steel stylet. After surgery, rats were allowed to recover for 3 and 5 days for i.c.v. and local surgeries, respectively, and were daily handled to adapt them to the infusion procedure, while stylets were cleaned. After completion of the experiment, i.c.v cannula placement was verified post mortem by infusion of 2-3 µl black ink (Pelikan 4001, Hannover, Germany). Local cannula placement was histologically confirmed with Nissel staining.

Immunohistochemistry and Microscopical analyses

Experimental animals were deeply anesthetized (ketamine/xylazine 3:1), transcardially perfused with 0.1M phosphate-buffered saline (PBS) followed by 4% paraformaldehyde. Fixated brains were removed and post-fixated for 1h (4% paraformaldehyde), submerged in sucrose solution (30%), snap-frozen on dry ice, and cut into 40-μm thick sections for immunohistochemistry.

Fos-staining started by washing the sections 3 x 10 min with fresh 0.1M PBS, followed by 30 min of 3% $H_2O_2$ and another round of 3 x 10 min 0.1M PBS. Tissue was then pre-incubated for 30 min with 0.1 M PBS containing 5% normal goat serum and 0.3% Triton-X-100, and incubated overnight at 4°C with rabbit-anti-c-fos antibody (1:10.000, sc-52 lot F-0315, Santa Cruz Biotechnology Inc, Santa Cruz, CA, USA) in 0.1 M PBS containing 0.5% normal goat serum and 0.3% Triton-X-100 (PBS-BT). The next day, tissue was washed 3 x 20 min with 0.1 M PBS. Brains were then incubated for 90 min with PBS-T containing biotinylated goat-anti-rabbit antibody (1:800, BA-1000, Vector Laboratories, Inc., Burlingame, CA, USA), followed by 3 x 20 min washing with 0.1 M PBS. Staining was enhanced by 90 min of incubation with PBS-T containing ABC-vector (1:800, PK-6100, Vector Laboratories, Inc., Burlingame, CA, USA), followed by 3 x 20 min washing with 0.1 M PBS. The tissue was then stained with a DAB-staining kit (SK-4100, Vector Laboratories, Inc., Burlingame, CA, USA) resulting in a blue-black nuclear staining, and washed 3 x 15 min in 0.1 M PBS. Brain sections were mounted on adhesive microscope slides (Superfrost® Plus, Thermo Fisher Scientific Inc., Waltham, MA, USA), dehydrated in increasing ethanol concentrations, cleared and embedded in Roti®-Histol and Roti®-Histokitt (Carl Roth GmbH, Karslruhe, Germany), and coverslipped.

Single labeling of Fos was analyzed in ten fore- and midbrain areas as derived from the Paxinos rat brain atlas[34]: the anterior nucleus accumbens core and shell, anterior cingulate cortex 1, prelimbic and infralimbic cortex and anterior agranular insular cortex (at or around Bregma 2.70); the posterior nucleus accumbens core and shell, posterior cingulate cortex 1 and 2, posterior agranular insular cortex (pAIC) (at or around Bregma 1.60); the dorsal and ventral lateral septum and medial preoptic area (at or around Bregma -0.30); the basolateral amygdala and lateral hypothalamic area (at or around Bregma -2.12); the central amygdala and anterior hypothalamic area (at or around Bregma -2.80); and the posterodorsal medial amygdala, ventrolateral ventromedial hypothalamus and lateral habenula (at or around

Bregma -3.20). Brain areas were selected based on their known role in aggression, consensual mating, and/or social decision-making[35–40].

Brain sections were imaged using a Leica DM5000B microscope connected to a personal computer; photographs were taken at 10x magnification using Leica Application Suite v3.7 software. Images were analyzed using Fiji ImageJ software[41]. The number of Fos-immunoreactive neurons was counted in a 200 x 200μm square placed over a representative area within the selected nucleus by an experienced analyst blinded to the experimental conditions.
